## Supplementary Information File for "An ambiguous N-terminus drives the dual targeting of an antioxidant protein Thioredoxin peroxidase (TgTPx1/2) to endosymbiotic organelles in *Toxoplasma gondii*"

**Supplementary Figures**

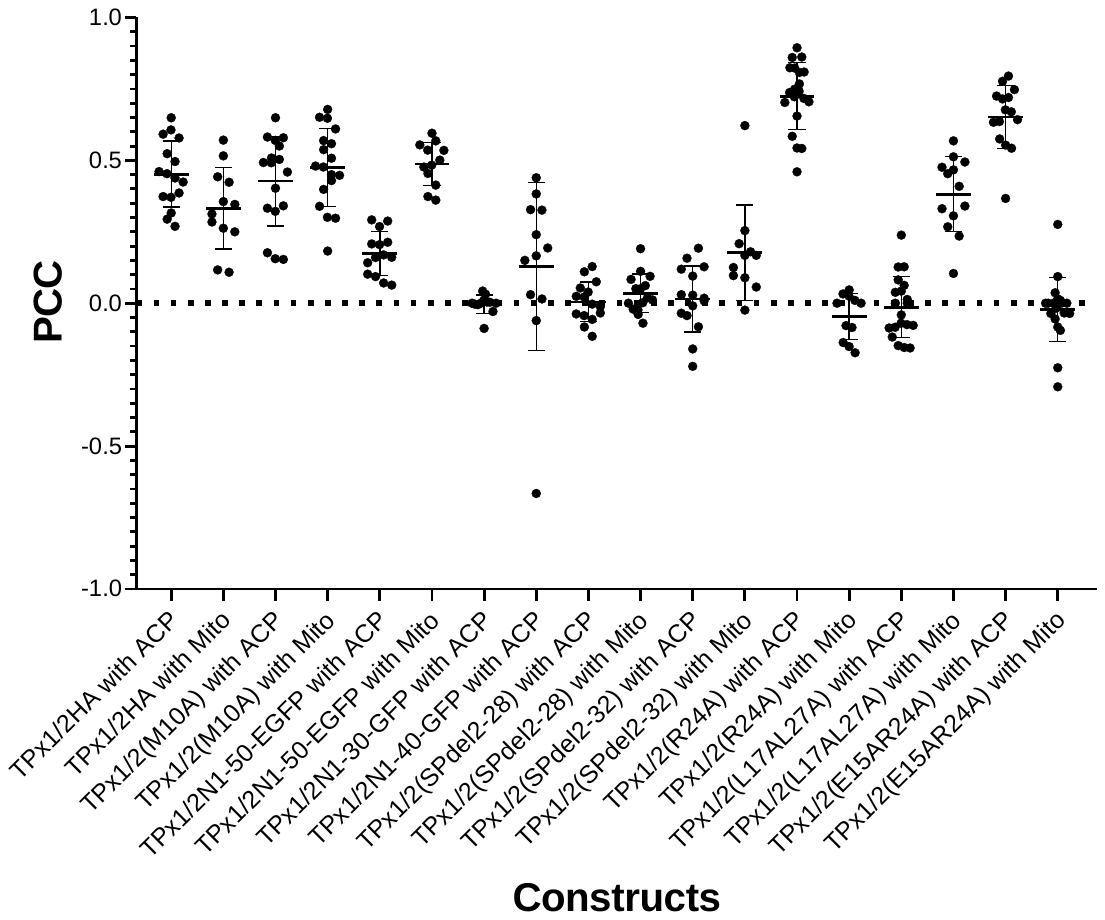
**Supplementary figure S1**: A dot plot of the Pearson’s correlation coefficients (PCC) for colocalization between the proteins expressed by the individual constructs the appropriate markers used in this study. Here, ACP is used as an apicoplast marker protein while SP‑TP‑SOD2-GFP and Mitotracker Red are used as mitochondrial markers (Mito).

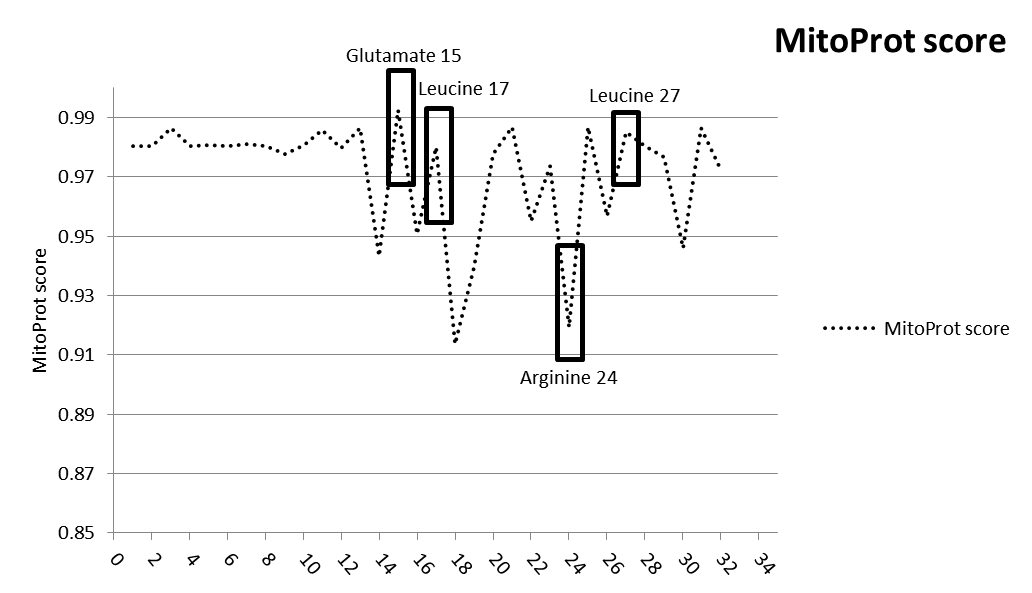

**Supplementary figure S2**: Determination of MitoProt scores of alanine scanning mutation analysis of the N-terminus of TgTPx1/2 employing the new version of MitoProt II. The full‑length TgTPx1/2 protein was used as an input sequence in the two bioinformatics software. For each of the first 30 amino acids, the native amino acid was replaced with alanine and the resulting full length proteins were used as input sequence. Default settings were employed for the analysis. The MitoProt (dashed black line) scores were plotted for the WT and the mutant proteins.

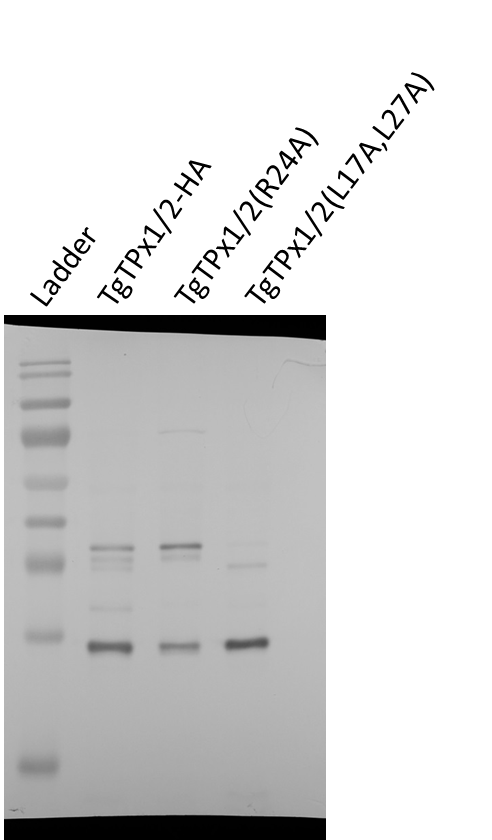

**S****upplementary figure S3**: Full-length Western blots displayed in main figures. High contrast images were not generated and hence multiple exposure images are not included here.
